## Supplemental Fig 1 and 2 for "Contribution of the *Tobamovirus* resistance gene *Tm-1* to control of ToBRFV resistance in tomato"

### Supplementary data

#### S1. *Tm-1* coding sequence nucleotide alignment

```
Tm-1 GCR237       1   ATGGCAACTGCACAGAGTAATTCTCCTCGAGTTTTCTGTATCGGAACAGCTGATACAAAATTCGACGAGC
Tm-1 VC554 1st    1   ATGGCAACTGCACAGAGTAATTCTCCTCGAGTTTTCTGTATCGGAACAGCTGATACAAAATTCGACGAGC
tm-1 GCR26        1   ATGGCAACTGCACAGAGTAATTCTCCTCGAGTTTTCTGTATCGGAACAGCTGATACTAAATTCGACGAGC
tm-1 Moneymaker   1   ATGGCAACTGCACAGAGTAATTCTCCTCGAGTTTTCTGTATCGGAACAGCTGATACTAAATTCGACGAGC
tm-1 VC532        1   ATGGCAACTGCACAGAGTAATTCTCCTCGAGTTTTCTGTATCGGAACAGCTGATACTAAATTCGACGAGC
Tm-1 VC554 2nd    1   ATGGCAAGTGCACAGAGTAATTCTCCTCGAGTTTTCTGTATTGGAACAGCTGATACTAAATTCGACGAGC

Tm-1 GCR237       71  TTCGTTTCCTCTCCGAGCATGTGAGATCCAGTCTTAACAGCTTCTCCAATAAATCCTCATTCAAGGTAGG
Tm-1 VC554 1st    71  TTCGTTTCCTCTCCGAGCATGTGAGATCCAGTCTTAACAGCTTCTCCAATAAATCCTCATTCAAGGTAGG
tm-1 GCR26        71  TTCGTTTCCTCTCCGAGCATGTGAGATCCAGTCTTAACAGCTTCTCCAATAAATCCTCATTCAAGGTAGG
tm-1 Moneymaker   71  TTCGTTTCCTCTCCGAGCATGTGAGATCCAGTCTTAACAGCTTCTCCAATAAATCCTCATTCAAGGTAGG
tm-1 VC532        71  TTCGTTTCCTCTCCGAGCATGTGAGATCCAGTCTTAACAGCTTCTCCAATAAATCCTCATTCAAGGTAGG
Tm-1 VC554 2nd    71  TTCGTTTCCTCTCCCAATATGTGAGATCCAGTCTTAACAGCTTCTCCAATAAATCCTCATTCAAGGTCGG

Tm-1 GCR237       141 AGTGACAGTTGTTGATGTCTCAACCAGCTGGAAGGAGACAAATAGTTGTGCTGATTTTGATTTTGTACCG
Tm-1 VC554 1st    141 AGTGACAGTTGTTGATGTCTCAACCAGCTGGAAGGAGACAAATAGTTGTGCTGATTTTGATTTTGTACCG
tm-1 GCR26        141 AGTGACAGTTGTTGATGTCTCAACCAGCCGGAAGGAGACAAATAGTTGTGCTGATTTTGATTTTGTACCG
tm-1 Moneymaker   141 AGTGACAGTTGTTGATGTCTCAACCAGCCGGAAGGAGACAAATAGTTGTGCTGATTTTGATTTTGTACCG
tm-1 VC532        141 AGTGACAGTTGTTGATGTCTCAACCAGCCGGAAGGAGACAAATAGTTGTGCTGATTTTGATTTTGTACCG
Tm-1 VC554 2nd    141 AGTCACAGTTGTTGATGTCTCAACCAGCCTAAAGGAGACAAATGGTTGTGCTGATTTTGATTTTGTGCCG

Tm-1 GCR237       211 AGTAAGGATGTGCTGTCATGCCATACACTAGGGGAAGAAACTATGGGCACGTTTGCAGATATTAGAGGCC
Tm-1 VC554 1st    211 AGTAAGGATGTGCTGTCATGCCATACACTAGGGGAAGAAACTATGGGCACGTTTGCAGATATTAGAGGCC
tm-1 GCR26        211 AGTAAGGATGTGCTGTCATGCTATGCACGAGGGGAAGGAACTGTGGGCAGGTTTCCAGATATTAGAGGCC
tm-1 Moneymaker   211 AGTAAGGATGTGCTGTCATGCTATGCACGAGGGGAAGGAACTGTGGGCAGGTTTCCAGATATTAGAGGCC
tm-1 VC532        211 AGTAAGGATGTGCTGTCATGCTATGCACGAGGGGAAGGAACTGTGGGCAGGTTTCCAGATATTAGAGGCC
Tm-1 VC554 2nd    211 AGGAAGGATGTGCTGTCCTGCTATGCACAAGGGGGAGAATCTGTGGTCCAGCTTCCAGATGATAGAGGCC

Tm-1 GCR237       281 TAGCTATTGCAATCATGAGCAAAGCTCTTGAAACTTTCCTAAGTATAGCTAATGATGAACAGAATCTTGC
Tm-1 VC554 1st    281 TAGCTATTGCAATCATGAGCAAAGCTCTTGAAACTTTCCTAAGTATAGCTAATGATGAACAGAATCTTGC
tm-1 GCR26        281 AAGCTATTGCAATCATGAACAAAGCTCTGGAAACTTTCCTAAGTAAAGCTAATGGTGAACAGAATCTTGC
tm-1 Moneymaker   281 AAGCTATTGCAATCATGAACAAAGCTCTGGAAACTTTCCTAAGTAAAGCTAATGGTGAACAGAATCTTGC
tm-1 VC532        281 AAGCTATTGCAATCATGAACAAAGCTCTGGAAACTTTCCTAAGTAAAGCTAATGGTGAACAGAATCTTGC
Tm-1 VC554 2nd    281 AAGCTATTGCAATCATGAACAAAGCTTTTCAAACTTTCCTAAGCAAAGCTAATGGTGAACAGAATCTTGC

Tm-1 GCR237       351 TGGAGTAATTGGCCTTGGGGGTAGTGGAGGAACATCTCTATTGTCATCTGCCTTCCGATCTCTTCCAATT
Tm-1 VC554 1st    351 TGGAGTAATTGGCCTTGGGGGTAGTGGAGGAACATCTCTATTGTCATCTGCCTTCCGATCTCTTCCAATT
tm-1 GCR26        351 TGGAGTGATTGGCCTTGGGGGTAGTGGAGGAACATCTCTATTGTCATCTGCCTTCCGATCTCTTCCAATT
tm-1 Moneymaker   351 TGGAGTGATTGGCCTTGGGGGTAGTGGAGGAACATCTCTATTGTCATCTGCCTTCCGATCTCTTCCAATT
tm-1 VC532        351 TGGAGTGATTGGCCTTGGGGGTAGTGGAGGAACATCTCTATTGTCATCTGCCTTCCGATCTCTTCCAATT
Tm-1 VC554 2nd    351 TGGAGTGATTGGCCTTGGGGGTAGTGGAGGAACATCTCTATTGTCATCTGCCTTCCGATCTCTTCCAATT
```

```
Tm-1 GCR237       421 GGGATCCCAAAAGTTATAATATCTACAGTTGCCAGTGGTCAAACTGAATCTTATATTGGAACATCAGACT
Tm-1 VC554 1st    421 GGGATCCCAAAAGTTATAATATCTACAGTTGCCAGTGGTCAAACTGAATCTTATATTGGAACATCAGACT
tm-1 GCR26        421 GGGATCCCAAAAGTTATAATATCTACAGTTGCCAGTGGCCAAACTGAATCTTATATTGGAACATCAGACT
tm-1 Moneymaker   421 GGGATCCCAAAAGTTATAATATCTACAGTTGCCAGTGGCCAAACTGAATCTTATATTGGAACATCAGACT
tm-1 VC532        421 GGGATCCCAAAAGTTATAATATCTACAGTTGCCAGTGGCCAAACTGAATCTTATATTGGAACATCAGACT
Tm-1 VC554 2nd    421 GGAATCCCAAAAGTTATAATATCTACAGTTGCCAGTGGTCAAACTGAATCTTATATTGGAACATCAGACT

Tm-1 GCR237       491 TGGTATTGTTTCCTTCAGTTGTAGATATTTGTGGGATTAACAATGTCAGTAAGGTTGTTCTATCTAATGC
Tm-1 VC554 1st    491 TGGTATTGTTTCCTTCAGTTGTAGATATTTGTGGGATTAACAATGTCAGTAAGGTTGTTCTATCTAATGC
tm-1 GCR26        491 TGGTATTGTTTCCTTCAGTTGTAGATATTTGTGGGATTAACAATGTTAGTAAGGTTGTTCTATCTAATGC
tm-1 Moneymaker   491 TGGTATTGTTTCCTTCAGTTGTAGATATTTGTGGGATTAACAATGTTAGTAAGGTTGTTCTATCTAATGC
tm-1 VC532        491 TGGTATTGTTTCCTTCAGTTGTAGATATTTGTGGGATTAACAATGTTAGTAAGGTTGTTCTATCTAATGC
Tm-1 VC554 2nd    491 TGGTATTGTTTCCTTCAGTTGTAGATATTTGTGGGATTAACAATGTTAGTAAGGTTATTCTATCTAATGC

Tm-1 GCR237       561 GGGTGCAGCATTTGCTGGAATGGTGATCGGGAGGCTTGAAAGTTCAAAAGAGCATAGCATCACTAATGGA
Tm-1 VC554 1st    561 GGGTGCAGCATTTGCTGGAATGGTGATCGGGAGGCTTGAAAGTTCAAAAGAGCATAGCATCACTAATGGA
tm-1 GCR26        561 GGGTGCAGCATTTGCTGGAATGGTGATTGGAAGGCTTGAAAGTTCAAAAGAGCATAGCATCACTAATGGA
tm-1 Moneymaker   561 GGGTGCAGCATTTGCTGGAATGGTGATTGGAAGGCTTGAAAGTTCAAAAGAGCATAGCATCACTAATGGA
tm-1 VC532        561 GGGTGCAGCATTTGCTGGAATGGTGATTGGAAGGCTTGAAAGTTCAAAAGAGCATAGCATCACTAATGGA
Tm-1 VC554 2nd    561 GGGTGCAGCATTTGCTGGAATGGTGATCGGAAGGCTTGAAACTTCAAAAGAGAATAGCATCACTACTGGA

Tm-1 GCR237       631 AAGTTTACAGTTGGTGTAACTATGTTTGGGGTTACGACTCCTTGTGTTAATGCTGTCAAAGAAAGATTAG
Tm-1 VC554 1st    631 AAGTTTACAGTTGGTGTAACTATGTTTGGGGTTACGACTCCTTGTGTTAATGCTGTCAAAGAAAGATTAG
tm-1 GCR26        631 AAGTTTACAGTTGGTGTAACTATGTTTGGGGTTACGACTCCTTGTGTTAATGCTGTCAAAGAAAGATTAG
tm-1 Moneymaker   631 AAGTTTACAGTTGGTGTAACTATGTTTGGGGTTACGACTCCTTGTGTTAATGCTGTCAAAGAAAGATTAG
tm-1 VC532        631 AAGTTTACAGTTGGTGTAACTATGTTTGGGGTTACGACTCCTTGTGTTAATGCTGTCAAAGAAAGATTAG
Tm-1 VC554 2nd    631 AAGTTTACAGTTGGTGTAACTATGTTTGGGGTTACGACTCCTTGTGTTAATGCTGTCAAAGAAAGATTAG

Tm-1 GCR237       701 TGAAAGAAGGATATGAGACTTTGGTGTTCCATGCCACGGGTGTCGGGGGCAGGGCCATGGAGGATCTTGT
Tm-1 VC554 1st    701 TGAAAGAAGGATATGAGACTTTGGTGTTCCATGCCACGGGTGTCGGGGGCAGGGCCATGGAGGATCTTGT
tm-1 GCR26        701 TGAAAGAAGGATATGAGACTTTGGTGTTCCATGCCACGGGTGTCGGGGGCAGGGCCATGGAGGATCTTGT
tm-1 Moneymaker   701 TGAAAGAAGGATATGAGACTTTGGTGTTCCATGCCACGGGTGTCGGGGGCAGGGCCATGGAGGATCTTGT
tm-1 VC532        701 TGAAAGAAGGATATGAGACTTTGGTGTTCCATGCCACGGGTGTCGGGGGCAGGGCCATGGAGGATCTTGT
Tm-1 VC554 2nd    701 TGAAAGAAGGATATGAGACTTTGGTTTTCCATGCCACGGGTGTCGGGGGCAGGGCCATGGAGGATCTTGT

Tm-1 GCR237       771 TAGAGGAGGTTTTATACAGGGTGTGCTGGATATTACGACAACTGAGGTTGCAGATTACGTAGTTGGAGGA
Tm-1 VC554 1st    771 TAGAGGAGGTTTTATACAGGGTGTGCTGGATATTACGACAACTGAGGTTGCAGATTACGTAGTTGGAGGA
tm-1 GCR26        771 TAGAGGAGGTTTTATACAGGGTGTGCTGGATATTACGACAACTGAGGTTGCAGATTACGTAGTTGGAGGA
tm-1 Moneymaker   771 TAGAGGAGGTTTTATACAGGGTGTGCTGGATATTACGACAACTGAGGTTGCAGATTACGTAGTTGGAGGA
tm-1 VC532        771 TAGAGGAGGTTTTATACAGGGTGTGCTGGATATTACGACAACTGAGGTTGCAGATTACGTAGTTGGAGGA
Tm-1 VC554 2nd    771 TAGAGCAGGTTTTATACAGGGCGTGCTGGATATTACGACAACTGAGGTTGCAGATTACGTAGTTGGAGGA

Tm-1 GCR237       841 GTAATGGCATGTGATAGTTCCCGATTTGATGCAATATTAGAGAAGAAAATTCCTTTGGTTCTGAGTGTGG
Tm-1 VC554 1st    841 GTAATGGCATGTGATAGTTCCCGATTTGATGCAATATTAGAGAAGAAAATTCCTTTGGTTCTGAGTGTGG
tm-1 GCR26        841 GTAATGGCATGTGATAGTTCCCGATTTGATGCAATATTAGAGAAGAAAATTCCTTTGGTTCTGAGTGTGG
tm-1 Moneymaker   841 GTAATGGCATGTGATAGTTCCCGATTTGATGCAATATTAGAGAAGAAAATTCCTTTGGTTCTGAGTGTGG
tm-1 VC532        841 GTAATGGCATGTGATAGTTCCCGATTTGATGCAATATTAGAGAAGAAAATTCCTTTGGTTCTGAGTGTGG
Tm-1 VC554 2nd    841 GTAATGGCATGTGATAGTTCCCGATTTGATGCAATATTAGAGAAGAAAATTCCTTTGGTTCTGAGTGTGG
```

| | | |
|---|---|---|
| ***Tm-1 GCR237*** | 911 | GAGCACTGGATATGGTGAATTTTGGTCCTAAAACTACCATACCTCCTGAGTTTCAACAAAGAAAGATCCA |
| ***Tm-1 VC554 1st*** | 911 | GAGCACTGGATATGGTGAATTTTGGTCCTAAAACTACCATACCTCCTGAGTTTCAACAAAGAAAGATCCA |
| ***tm-1 GCR26*** | 911 | GAGCACTGGATATGGTGAATTTTGGTCCTAAAACTACCATACCTCCTGAGTTTCAGCAAAGAAAGATTCA |
| ***tm-1 Moneymaker*** | 911 | GAGCACTGGATATGGTGAATTTTGGTCCTAAAACTACCATACCTCCTGAGTTTCAGCAAAGAAAGATTCA |
| ***tm-1 VC532*** | 911 | GAGCACTGGATATGGTGAATTTTGGTCCTAAAACTACCATACCTCCTGAGTTTCAGCAAAGAAAGATTCA |
| ***Tm-1 VC554 2nd*** | 911 | GAGCACTGGATATGGTGAATTTTGGTCCTAAAACTACCATACCACCTGAGTTTCAGCAAAGAAAGATTCA |
| ***Tm-1 GCR237*** | 981 | TGAACATAATGAGCAGGTTTCCCTAATGCGTACTACAGTAGGTGAAAATAAGAAATTTGCTGCATTTATA |
| ***Tm-1 VC554 1st*** | 981 | TGAACATAATGAGCAGGTTTCCCTAATGCGTACTACAGTAGGTGAAAATAAGAAATTTGCTGCATTTATA |
| ***tm-1 GCR26*** | 981 | TCAACATAATGAGCAGGTTTCCCTAATGCATACTACAGTAGGTGAAAATAAGAAATTTGCTGCATTTATA |
| ***tm-1 Moneymaker*** | 981 | TCAACATAATGAGCAGGTTTCCCTAATGCATACTACAGTAGGTGAAAATAAGAAATTTGCTGCATTTATA |
| ***tm-1 VC532*** | 981 | TCAACATAATGAGCAGGTTTCCCTAATGCGTACTACAGTAGGTGAAAATAAGAAATTTGCTGCATTTATA |
| ***Tm-1 VC554 2nd*** | 981 | TCAACATAATGAGCAGGTTTCCATAATGCGTACTACAGTAGGTGAAAATAAGAAATTTGCTGCATTTATA |
| ***Tm-1 GCR237*** | 1051 | GCAGAAAAGTTGAACAAGGCATCATCAAGTGTATGTGTTTGCTTGCCAGAGAAAGGCGTGTCTGCATTGG |
| ***Tm-1 VC554 1st*** | 1051 | GCAGAAAAGTTGAACAAGGCATCATCAAGTGTATGTGTTTGCTTGCCAGAGAAAGGCGTGTCTGCATTGG |
| ***tm-1 GCR26*** | 1051 | GCAGAAAAGTTGAACAAGGCATCATCAAGTGTATGTGTTTGCTTGCCAGAGAAAGGCGTGTCTGCATTGG |
| ***tm-1 Moneymaker*** | 1051 | GCAGAAAAGTTGAACAAGGCATCATCAAGTGTATGTGTTTGCTTGCCAGAGAAAGGCGTGTCTGCATTGG |
| ***tm-1 VC532*** | 1051 | GCAGAAAAGTTGAACAAGGCATCATCAAGTGTATGTGTTTGCTTGCCAGAGAAAGGCGTGTCTGCATTGG |
| ***Tm-1 VC554 2nd*** | 1051 | GCTGAAAAGTTGAACAAGGCATCATCAAGTGTATGTGTTTGCTTGCCAGAGAAAGGTGTGTCTGCATTGG |
| ***Tm-1 GCR237*** | 1121 | ATGCACCCGGGAAAGACTTTTATGATCCTGAGGCAACTAGTTGTCTTACACGTGAACTACAGATGCTTCT |
| ***Tm-1 VC554 1st*** | 1121 | ATGCACCCGGGAAAGACTTTTATGATCCTGAGGCAACTAGTTGTCTTACACGTGAACTACAGATGCTTCT |
| ***tm-1 GCR26*** | 1121 | ATGCACCCGGGAAAGACTTTTATGATCCTGAGGCAACTAGTTGTCTTACACATGAACTACAGATGCTTCT |
| ***tm-1 Moneymaker*** | 1121 | ATGCACCCGGGAAAGACTTTTATGATCCTGAGGCAACTAGTTGTCTTACACATGAACTACAGATGCTTCT |
| ***tm-1 VC532*** | 1121 | ATGCACCCGGGAAAGACTTTTATGATCCTGAGGCAACTAGTTGTCTTACACATGAACTACAGATGCTTCT |
| ***Tm-1 VC554 2nd*** | 1121 | ATGCACCGGGGAAAGAATTTTATGATCCTGAGGCAACTAGTTGTCTTACACATGAGCTACTGATGCTTCT |
| ***Tm-1 GCR237*** | 1191 | TGAAAATAATGAACGTTGTCAGGTTAAGGTCCTCCCTTACCATATCAATGATGCGGAGTTTGCAAATGCT |
| ***Tm-1 VC554 1st*** | 1191 | TGAAAATAATGAACGTTGTCAGGTTAAGGTCCTCCCTTACCATATCAATGATGCGGAGTTTGCAAATGCT |
| ***tm-1 GCR26*** | 1191 | TGAAAATAATGAACGTTGTCAGGTTAAGGTCTACCCTTACCATATCAATGATGTGGAGTTTGCAAATGCT |
| ***tm-1 Moneymaker*** | 1191 | TGAAAATAATGAACGTTGTCAGGTTAAGGTCTACCCTTACCATATCAATGATGTGGAGTTTGCAAATGCT |
| ***tm-1 VC532*** | 1191 | TGAAAATAATGAACGTTGTCAGGTTAAGGTCTACCCTTACCATATCAATGATGTGGAGTTTGCAAATGCT |
| ***Tm-1 VC554 2nd*** | 1191 | TGAAAACAATGAACGTTGTCAGGTTAAGGTCTTCCCTTGCCATATCAATGATGCGGAGTTTGCAAATGCT |
| ***Tm-1 GCR237*** | 1261 | TTAGTTGATTCATTCTTGGAAATCTCTCCGAAATCTAGACACGTAGAATGTCAGCCAGCTGAGTCCAAAT |
| ***Tm-1 VC554 1st*** | 1261 | TTAGTTGATTCATTCTTGGAAATCTCTCCGAAATCTAGACACGTAGAATGTCAGCCAGCTGAGTCCAAAT |
| ***tm-1 GCR26*** | 1261 | TTAGTTGATTCATTTTTGGAAATGTCTCCGAAATCTGGACACGTAGAATGTCAGACAGCTGAGTCCAAAT |
| ***tm-1 Moneymaker*** | 1261 | TTAGTTGATTCATTTTTGGAAATGTCTCCGAAATCTGGACACGTAGAATGTCAGACAGCTGAGTCCAAAT |
| ***tm-1 VC532*** | 1261 | TTAGTTGATTCATTTTTGGAAATGTCTCCGAAATCTGGACACGTAGAATGTCAGACAGCTGAGTCCAAAT |
| ***Tm-1 VC554 2nd*** | 1261 | TTAGTTGATTCATTCTTGGAAGTCTCTCCGAAATCTAGACACGTAGAATGTCAGCCAGCTGAGTCCAAAT |
| ***Tm-1 GCR237*** | 1331 | CTATCCAAGACATTCAGAATGATAATGCTGTTCTAGAGAAATATCCCTCATGCAACGGGAAAAACTTTTC |
| ***Tm-1 VC554 1st*** | 1331 | CTATCCAAGACATTCAGAATGATAATGCTGTTCTAGAGAAATATCCCTCATGCAACGGGAAAAACTTTTC |
| ***tm-1 GCR26*** | 1331 | CTATACAAGGCATTCAGAATGTTAATGCTGTTCTAGAGAAATATCCCTCATGCAACGGGAAAAACTTTTC |
| ***tm-1 Moneymaker*** | 1331 | CTATACAAGGCATTCAGAATGTTAATGCTGTTCTAGAGAAATATCCCTCATGCAACGGGAAAAACTTTTC |
| ***tm-1 VC532*** | 1331 | CTATACAAGGCATTCAGAATGTTAATGCTGTTCTAGAGAAATATCCCTCATGCAACGGGAAAAACTTTTC |
| ***Tm-1 VC554 2nd*** | 1331 | GTATCCAAGACATTCAGAATGATAATGCTGTTCTAGAGAAATATCCCTCATGCAACGGGAAAAACTTTTC |

```
Tm-1 GCR237       1401 TCGCCTGAATGACTTTCCAAATGCAAAACCAGAAACTTTGCAGAAAAGAACTGTGATACTGCAGAAATTG
Tm-1 VC554 1st    1401 TCGCCTGAATGACTTTCCAAATGCAAAACCAGAAACTTTGCAGAAAAGAACTGTGATACTGCAGAAATTG
tm-1 GCR26        1401 TCGCCTGAATGACTTTCCAAATGCAAAACCAGAAACTTTGCAGAAAAGAATTGTGATACTGCAGAAATTG
tm-1 Moneymaker   1401 TCGCCTGAATGACTTTCCAAATGCAAAACCAGAAACTTTGCAGAAAAGAATTGTGATACTGCAGAAATTG
tm-1 VC532        1401 TCGCCTGAATGACTTTCCAAATGCAAAACCAGAAACTTTGCAGAAAAGAATTGTGATACTGCAGAAATTG
Tm-1 VC554 2nd    1401 TCGCCTGAATGACTTTCCAAATGCAAAACCAGAAACTTTGCAGAAAAGAACTGTGATACTGCAGAAATTG

Tm-1 GCR237       1471 AAAGATCAAATAAGTAAGGGCAAGCCTATTATTGGGGCTGGTGCTGGTACAGGTATTTCTGCTAAGTTTG
Tm-1 VC554 1st    1471 AAAGATCAAATAAGTAAGGGCAAGCCTATTATTGGGGCTGGTGCTGGTACAGGTATTTCTGCTAAGTTTG
tm-1 GCR26        1471 AAAGATCAAATAAGTAAGGGCAAGCCTATTATTGGGGCTGGTGCTGGTACAGGTATTTCTGCTAAGTTTG
tm-1 Moneymaker   1471 AAAGATCAAATAAGTAAGGGCAAGCCTATTATTGGGGCTGGTGCTGGTACAGGTATTTCTGCTAAGTTTG
tm-1 VC532        1471 AAAGATCAAATAAGTAAGGGCAAGCCTATTATTGGGGCTGGTGCTGGTACAGGTATTTCTGCTAAGTTTG
Tm-1 VC554 2nd    1471 AAAGATCAAATAAGTAAGGGCAAGCCTATTATTGGGGCTGGAGCTGGTACAGGTATTTCTGCTAAGTTTG

Tm-1 GCR237       1541 AGGAAGCTGGTGGTGTAGATTTGATTGTCTTGTACAACTCAGGGCGCTTTAGGATGGCAGGAAGGGGATC
Tm-1 VC554 1st    1541 AGGAAGCTGGTGGTGTAGATTTGATTGTCTTGTACAACTCAGGGCGCTTTAGGATGGCAGGAAGGGGATC
tm-1 GCR26        1541 AGGAAGCTGGTGGTGTAGATTTGATTGTCTTGTACAACTCAGGGCGCTTTAGGATGGCAGGAAGGGGATC
tm-1 Moneymaker   1541 AGGAAGCTGGTGGTGTAGATTTGATTGTCTTGTACAACTCAGGGCGCTTTAGGATGGCAGGAAGGGGATC
tm-1 VC532        1541 AGGAAGCTGGTGGTGTAGATTTGATTGTCTTGTACAACTCAGGGCGCTTTAGGATGGCAGGAAGGGGATC
Tm-1 VC554 2nd    1541 AGGAAGCTGGTGGTGTGGATTTGATTGTCTTGTACAACTCAGGGCGCTTTAGGATGGCAGGAAGGGGATC

Tm-1 GCR237       1611 CTTAGCTGGTCTACTGCCCTTTGCTGATGCAAATGCCATTGTACTTGAGATGGCCAACGAAGTATTGCCT
Tm-1 VC554 1st    1611 CTTAGCTGGTCTACTGCCCTTTGCTGATGCAAATGCCATTGTACTTGAGATGGCCAACGAAGTATTGCCT
tm-1 GCR26        1611 CTTAGCTGGTCTATTGCCCTTTGCTGATGCAAATGCCATTGTACTTGAGATGGCCAACGAAGTATTGCCT
tm-1 Moneymaker   1611 CTTAGCTGGTCTATTGCCCTTTGCTGATGCAAATGCCATTGTACTTGAGATGGCCAACGAAGTATTGCCT
tm-1 VC532        1611 CTTAGCTGGTCTATTGCCCTTTGCTGATGCAAATGCCATTGTACTTGAGATGGCCAACGAAGTATTGCCT
Tm-1 VC554 2nd    1611 CTTAGCTGGTCTATTGCCCTTTGCTGATGCAAATGCCATTGTACTTGAGATGGCCAACGAAGTATTGCCG

Tm-1 GCR237       1681 GTGGTTAAGGAAGTGGCAGTTCTGGCTGGAGTTTGTGCTACTGATCCTTTCCGCAGGATGGACAACTTCC
Tm-1 VC554 1st    1681 GTGGTTAAGGAAGTGGCAGTTCTGGCTGGAGTTTGTGCTACTGATCCTTTCCGCAGGATGGACAACTTCC
tm-1 GCR26        1681 GTGGTTAAGGAAGTGGCAGTTCTGGCTGGAGTTTGTGCTACTGATCCTTTCCGCAGGATGGACAACTTCC
tm-1 Moneymaker   1681 GTGGTTAAGGAAGTGGCAGTTCTGGCTGGAGTTTGTGCTACTGATCCTTTCCGCAGGATGGACAACTTCC
tm-1 VC532        1681 GTGGTTAAGGAAGTGGCAGTTCTGGCTGGAGTTTGTGCTACTGATCCTTTCCGCAGGATGGACAACTTCC
Tm-1 VC554 2nd    1681 GTGGTTAAGGAAGTGGCAGTTCTGGCTGGAGTTTGTGCAACTGATCCTTTCCGCAGGATGGACAACTTCC

Tm-1 GCR237       1751 TGAAGCAGTTGGAATCCGTTGGATTCTGTGGGGTGCAAAACTTTCCAACTGTTGGTCTGTTTGACGGTAA
Tm-1 VC554 1st    1751 TGAAGCAGTTGGAATCCGTTGGATTCTGTGGGGTGCAAAACTTTCCAACTGTTGGTCTGTTTGACGGTAA
tm-1 GCR26        1751 TGAAGCAGTTGGAATCTGTTGGATTCTGTGGGGTGCAAAACTTTCCAACTGTTGGTCTGTTTGACGGTAA
tm-1 Moneymaker   1751 TGAAGCAGTTGGAATCTGTTGGATTCTGTGGGGTGCAAAACTTTCCAACTGTTGGTCTGTTTGACGGTAA
tm-1 VC532        1751 TGAAGCAGTTGGAATCTGTTGGATTCTGTGGGGTGCAAAACTTTCCAACTGTTGGTCTGTTTGACGGTAA
Tm-1 VC554 2nd    1751 TGAAGCAGTTGGAATCCGTTGGATTCTGTGGGGTGCAAAACTTTCCAACTGTTGGTCTGTTTGACGGTAA

Tm-1 GCR237       1821 CTTCAGACAAAATTTGGAAGAGACTGGAATGGGTTATGGCTTGGAGGTTGAGATGATTGCAGCAGCTCAC
Tm-1 VC554 1st    1821 CTTCAGACAAAATTTGGAAGAGACTGGAATGGGTTATGGCTTGGAGGTTGAGATGATTGCAGCAGCTCAC
tm-1 GCR26        1821 CTTCAGACAAAATTTGGAAGAGACTGGAATGGGTTATGGCTTGGAGGTTGAGATGATTGCAACAGCTCAT
tm-1 Moneymaker   1821 CTTCAGACAAAATTTGGAAGAGACTGGAATGGGTTATGGCTTGGAGGTTGAGATGATTGCAACAGCTCAT
tm-1 VC532        1821 CTTCAGACAAAATTTGGAAGAGACTGGAATGGGTTATGGCTTGGAGGTTGAGATGATTGCAACAGCTCAT
Tm-1 VC554 2nd    1821 CTTCAGACAAAATTTGGAAGAGACTGGAATGGGTTATGGCTTGGAGGTTGAGATGATTGCAACAGCTCAC
```

| | | |
|---|---|---|
| ***Tm-1 GCR237*** | 1891 | AGGATGGGCCTTTTGACAACCCCATATGCTTTCTGCCCAGATGAAGCAGTTGCTATGGCAGAAGCTGGTG |
| ***Tm-1 VC554 1st*** | 1891 | AGGATGGGCCTTTTGACAACCCCATATGCTTTCTGCCCAGATGAAGCAGTTGCTATGGCAGAAGCTGGTG |
| ***tm-1 GCR26*** | 1891 | AGGATGGGCCTTTTGACAACCCCATATGCTTTCTGCCCAGATGAAGCAGTTGCTATGGCAGAAGCTGGTG |
| ***tm-1 Moneymaker*** | 1891 | AGGATGGGCCTTTTGACAACCCCATATGCTTTCTGCCCAGATGAAGCAGTTGCTATGGCAGAAGCTGGTG |
| ***tm-1 VC532*** | 1891 | AGGATGGGCCTTTTGACAACCCCATATGCTTTCTGCCCAGATGAAGCAGTTGCTATGGCAGAAGCTGGTG |
| ***Tm-1 VC554 2nd*** | 1891 | AGGATGGGCCTTTTGACAACCCCATATGCTTTCTGCCCAGATGAAGCAGTTGCTATGGCAGAAGCTGGTG |
| ***Tm-1 GCR237*** | 1961 | CCGACATCATAGTTGCTCATATGGGGCTTACAACATCTGGTTCAATTGGTGCAAAAACAGCCGTCTCATT |
| ***Tm-1 VC554 1st*** | 1961 | CCGACATCATAGTTGCTCATATGGGGCTTACAACATCTGGTTCAATTGGTGCAAAAACAGCCGTCTCATT |
| ***tm-1 GCR26*** | 1961 | CCGACATCATAGTTGCTCATATGGGGCTTACAACATCTGGTTCAATTGGTGCAAAAACAGCTGTATCATT |
| ***tm-1 Moneymaker*** | 1961 | CCGACATCATAGTTGCTCATATGGGGCTTACAACATCTGGTTCAATTGGTGCAAAAACAGCTGTATCATT |
| ***tm-1 VC532*** | 1961 | CCGACATCATAGTTGCTCATATGGGGCTTACAACATCTGGTTCAATTGGTGCAAAAACAGCTGTATCATT |
| ***Tm-1 VC554 2nd*** | 1961 | CCGACATCATAGTTGCTCATATGGGGCTTACAACATCTGGTTCAATTGGTGCAAAAACAGCTGTCTCATT |
| ***Tm-1 GCR237*** | 2031 | GGAGGAAAGTGTAACTTGCGTTCAAGCTATTGCAGATGCTACTCATAGGATATATCCTGATGCAATTGTG |
| ***Tm-1 VC554 1st*** | 2031 | GGAGGAAAGTGTAACTTGCGTTCAAGCTATTGCAGATGCTACTCATAGGATATATCCTGATGCAATTGTG |
| ***tm-1 GCR26*** | 2031 | GGAGGAAAGTGTAACTTGCGTCCAAGCTATTGCAGATGCTACTCATAGGATAAATCCTGATGCAATTGTG |
| ***tm-1 Moneymaker*** | 2031 | GGAGGAAAGTGTAACTTGCGTCCAAGCTATTGCAGATGCTACTCATAGGATAAATCCTGATGCAATTGTG |
| ***tm-1 VC532*** | 2031 | GGAGGAAAGTGTAACTTGCGTCCAAGCTATTGCAGATGCTACTCATAGGATAAATCCTGATGCAATTGTG |
| ***Tm-1 VC554 2nd*** | 2031 | GGAGGAAAGTGTAACTTGCGTCCAAGCTATTGCAGATGCTACTCATAGGATAAATCCTGATGCAATTGTG |
| ***Tm-1 GCR237*** | 2101 | CTCTGCCATGGAGGCCCTATATCTTCCCCTGAAGAAGCAGCATATGTACTGAAGAGAACCACAGGAGTTC |
| ***Tm-1 VC554 1st*** | 2101 | CTCTGCCATGGAGGCCCTATATCTTCCCCTGAAGAAGCAGCATATGTACTGAAGAGAACCACAGGAGTTC |
| ***tm-1 GCR26*** | 2101 | CTCTGCCATGGAGGCCCTATATCTTCCCCTGAAGAAGCAGCATATGTACTGAAGAGAACCACAGGAGTTC |
| ***tm-1 Moneymaker*** | 2101 | CTCTGCCATGGAGGCCCTATATCTTCCCCTGAAGAAGCAGCATATGTACTGAAGAGAACCACAGGAGTTC |
| ***tm-1 VC532*** | 2101 | CTCTGCCATGGAGGCCCTATATCTTCCCCTGAAGAAGCAGCATATGTACTGAAGAGAACCACAGGAGTTC |
| ***Tm-1 VC554 2nd*** | 2101 | CTCTGCCATGGAGGCCCTATATCTTCCCCTGAAGAAGCAGCATATGTACTGAAGAGAACCACAGGAGTTC |
| ***Tm-1 GCR237*** | 2171 | ATGGATTTTATGGCGCTTCAAGCATGGAAAGACTACCAGTTGAGCAAGCTATAACTGCAACTGTCCAGCA |
| ***Tm-1 VC554 1st*** | 2171 | ATGGATTTTATGGCGCTTCAAGCATGGAAAGACTACCAGTTGAGCAAGCTATAACTGCAACTGTCCAGCA |
| ***tm-1 GCR26*** | 2171 | ATGGATTTTATGGCGCTTCAAGCATGGAAAGACTACCAGTTGAGCAAGCTATAACTGCAACTGTCCAACA |
| ***tm-1 Moneymaker*** | 2171 | ATGGATTTTATGGCGCTTCAAGCATGGAAAGACTACCAGTTGAGCAAGCTATAACTGCAACTGTCCAACA |
| ***tm-1 VC532*** | 2171 | ATGGATTTTATGGCGCTTCAAGCATGGAAAGACTACCAGTTGAGCAAGCTATAACTGCAACTGTCCAACA |
| ***Tm-1 VC554 2nd*** | 2171 | ATGGATTTTATGGCGCTTCAAGCATGGAAAGACTACCAGTTGAGCAAGCTATAACTGCAACTGTCCAGCA |
| ***Tm-1 GCR237*** | 2241 | GTACAAGTCTATTTCTATGGAGTGA |
| ***Tm-1 VC554 1st*** | 2241 | GTACAAGTCTATTTCTATGGAGTGA |
| ***tm-1 GCR26*** | 2241 | GTACAAGTCTATATCTATGGAGTGA |
| ***tm-1 Moneymaker*** | 2241 | GTACAAGTCTATATCTATGGAGTGA |
| ***tm-1 VC532*** | 2241 | GTACAAGTCTATATCTATGGAGTGA |
| ***Tm-1 VC554 2nd*** | 2241 | GTACAAGTCTATTTCTATGGAGTGA |

#### S2. *Tm-1* amino acid sequence alignment

```
Tm-1 GCR237        1    MATAQSNSPRVFCIGTADTKFDELRFLSEHVRSSLNSFSNKSSFKVGVTVVDVSTSWKETNSCADFDFVP
Tm-1 VC554 1st     1    MATAQSNSPRVFCIGTADTKFDELRFLSEHVRSSLNSFSNKSSFKVGVTVVDVSTSWKETNSCADFDFVP
tm-1 GCR26        1    MATAQSNSPRVFCIGTADTKFDELRFLSEHVRSSLNSFSNKSSFKVGVTVVDVSTSRKETNSCADFDFVP
tm-1 Moneymaker   1    MATAQSNSPRVFCIGTADTKFDELRFLSEHVRSSLNSFSNKSSFKVGVTVVDVSTSRKETNSCADFDFVP
tm-1 VC532        1    MATAQSNSPRVFCIGTADTKFDELRFLSEHVRSSLNSFSNKSSFKVGVTVVDVSTSRKETNSCADFDFVP
Tm-1 VC554 2nd     1    MASAQSNSPRVFCIGTADTKFDELRFLSQYVRSSLNSFSNKSSFKVGVTVVDVSTSLKETNGCADFDFVP

Tm-1 GCR237        71   SKDVLSCHTLGEETMGTFADIRGLAIAIMSKALETFLSIANDEQNLAGVIGLGGSGGTSLLSSAFRSLPI
Tm-1 VC554 1st     71   SKDVLSCHTLGEETMGTFADIRGLAIAIMSKALETFLSIANDEQNLAGVIGLGGSGGTSLLSSAFRSLPI
tm-1 GCR26        71   SKDVLSCYARGEGTVGRFPDIRGQAIAIMNKALETFLSKANGEQNLAGVIGLGGSGGTSLLSSAFRSLPI
tm-1 Moneymaker   71   SKDVLSCYARGEGTVGRFPDIRGQAIAIMNKALETFLSKANGEQNLAGVIGLGGSGGTSLLSSAFRSLPI
tm-1 VC532        71   SKDVLSCYARGEGTVGRFPDIRGQAIAIMNKALETFLSKANGEQNLAGVIGLGGSGGTSLLSSAFRSLPI
Tm-1 VC554 2nd     71   RKDVLSCYAQGGESVVQLPDDRGQAIAIMNKAFQTFLSKANGEQNLAGVIGLGGSGGTSLLSSAFRSLPI

Tm-1 GCR237        141  GIPKVIISTVASGQTESYIGTSDLVLFPSVVDICGINNVSKVVLSNAGAAFAGMVIGRLESSKEHSITNG
Tm-1 VC554 1st     141  GIPKVIISTVASGQTESYIGTSDLVLFPSVVDICGINNVSKVVLSNAGAAFAGMVIGRLESSKEHSITNG
tm-1 GCR26        141  GIPKVIISTVASGQTESYIGTSDLVLFPSVVDICGINNVSKVVLSNAGAAFAGMVIGRLESSKEHSITNG
tm-1 Moneymaker   141  GIPKVIISTVASGQTESYIGTSDLVLFPSVVDICGINNVSKVVLSNAGAAFAGMVIGRLESSKEHSITNG
tm-1 VC532        141  GIPKVIISTVASGQTESYIGTSDLVLFPSVVDICGINNVSKVVLSNAGAAFAGMVIGRLESSKEHSITNG
Tm-1 VC554 2nd     141  GIPKVIISTVASGQTESYIGTSDLVLFPSVVDICGINNVSKVILSNAGAAFAGMVIGRLETSKENSITTG

Tm-1 GCR237        211  KFTVGVTMFGVTTPCVNAVKERLVKEGYETLVFHATGVGGRAMEDLVRGGFIQGVLDITTTEVADYVVGG
Tm-1 VC554 1st     211  KFTVGVTMFGVTTPCVNAVKERLVKEGYETLVFHATGVGGRAMEDLVRGGFIQGVLDITTTEVADYVVGG
tm-1 GCR26        211  KFTVGVTMFGVTTPCVNAVKERLVKEGYETLVFHATGVGGRAMEDLVRGGFIQGVLDITTTEVADYVVGG
tm-1 Moneymaker   211  KFTVGVTMFGVTTPCVNAVKERLVKEGYETLVFHATGVGGRAMEDLVRGGFIQGVLDITTTEVADYVVGG
tm-1 VC532        211  KFTVGVTMFGVTTPCVNAVKERLVKEGYETLVFHATGVGGRAMEDLVRGGFIQGVLDITTTEVADYVVGG
Tm-1 VC554 2nd     211  KFTVGVTMFGVTTPCVNAVKERLVKEGYETLVFHATGVGGRAMEDLVRAGFIQGVLDITTTEVADYVVGG

Tm-1 GCR237        281  VMACDSSRFDAILEKKIPLVLSVGALDMVNFGPKTTIPPEFQQRKIHEHNEQVSLMRTTVGENKKFAAFI
Tm-1 VC554 1st     281  VMACDSSRFDAILEKKIPLVLSVGALDMVNFGPKTTIPPEFQQRKIHEHNEQVSLMRTTVGENKKFAAFI
tm-1 GCR26        281  VMACDSSRFDAILEKKIPLVLSVGALDMVNFGPKTTIPPEFQQRKIHQHNEQVSLMHTTVGENKKFAAFI
tm-1 Moneymaker   281  VMACDSSRFDAILEKKIPLVLSVGALDMVNFGPKTTIPPEFQQRKIHQHNEQVSLMHTTVGENKKFAAFI
tm-1 VC532        281  VMACDSSRFDAILEKKIPLVLSVGALDMVNFGPKTTIPPEFQQRKIHQHNEQVSLMRTTVGENKKFAAFI
Tm-1 VC554 2nd     281  VMACDSSRFDAILEKKIPLVLSVGALDMVNFGPKTTIPPEFQQRKIHQHNEQVSIMRTTVGENKKFAAFI

Tm-1 GCR237        351  AEKLNKASSSVCVCLPEKGVSALDAPGKDFYDPEATSCLTRELQMLLENNERCQVKVLPYHINDAEFANA
Tm-1 VC554 1st     351  AEKLNKASSSVCVCLPEKGVSALDAPGKDFYDPEATSCLTRELQMLLENNERCQVKVLPYHINDAEFANA
tm-1 GCR26        351  AEKLNKASSSVCVCLPEKGVSALDAPGKDFYDPEATSCLTHELQMLLENNERCQVKVYPYHINDVEFANA
tm-1 Moneymaker   351  AEKLNKASSSVCVCLPEKGVSALDAPGKDFYDPEATSCLTHELQMLLENNERCQVKVYPYHINDVEFANA
tm-1 VC532        351  AEKLNKASSSVCVCLPEKGVSALDAPGKDFYDPEATSCLTHELQMLLENNERCQVKVYPYHINDVEFANA
Tm-1 VC554 2nd     351  AEKLNKASSSVCVCLPEKGVSALDAPGKEFYDPEATSCLTHELLMLLENNERCQVKVFPCHINDAEFANA

Tm-1 GCR237        421  LVDSFLEISPKSRHVECQPAESKSIQDIQNDNAVLEKYPSCNGKNFSRLNDFPNAKPETLQKRTVILQKL
Tm-1 VC554 1st     421  LVDSFLEISPKSRHVECQPAESKSIQDIQNDNAVLEKYPSCNGKNFSRLNDFPNAKPETLQKRTVILQKL
tm-1 GCR26        421  LVDSFLEMSPKSGHVECQTAESKSIQGIQNVNAVLEKYPSCNGKNFSRLNDFPNAKPETLQKRIVILQKL
tm-1 Moneymaker   421  LVDSFLEMSPKSGHVECQTAESKSIQGIQNVNAVLEKYPSCNGKNFSRLNDFPNAKPETLQKRIVILQKL
tm-1 VC532        421  LVDSFLEMSPKSGHVECQTAESKSIQGIQNVNAVLEKYPSCNGKNFSRLNDFPNAKPETLQKRIVILQKL
Tm-1 VC554 2nd     421  LVDSFLEVSPKSRHVECQPAESKCIQDIQNDNAVLEKYPSCNGKNFSRLNDFPNAKPETLQKRTVILQKL
```

| | | |
|---|---|---|
| ***Tm-1 GCR237*** | 491 | KDQISKGKPIIGAGAGTGISAKFEEAGGVDLIVLYNSGRFRMAGRGSLAGLLPFADANAIVLEMANEVLP |
| ***Tm-1 VC554 1st*** | 491 | KDQISKGKPIIGAGAGTGISAKFEEAGGVDLIVLYNSGRFRMAGRGSLAGLLPFADANAIVLEMANEVLP |
| ***tm-1 GCR26*** | 491 | KDQISKGKPIIGAGAGTGISAKFEEAGGVDLIVLYNSGRFRMAGRGSLAGLLPFADANAIVLEMANEVLP |
| ***tm-1 Moneymaker*** | 491 | KDQISKGKPIIGAGAGTGISAKFEEAGGVDLIVLYNSGRFRMAGRGSLAGLLPFADANAIVLEMANEVLP |
| ***tm-1 VC532*** | 491 | KDQISKGKPIIGAGAGTGISAKFEEAGGVDLIVLYNSGRFRMAGRGSLAGLLPFADANAIVLEMANEVLP |
| ***Tm-1 VC554 2nd*** | 491 | KDQISKGKPIIGAGAGTGISAKFEEAGGVDLIVLYNSGRFRMAGRGSLAGLLPFADANAIVLEMANEVLP |
| ***Tm-1 GCR237*** | 561 | VVKEVAVLAGVCATDPFRRMDNFLKQLESVGFCGVQNFPTVGLFDGNFRQNLEETGMGYGLEVEMIAAAH |
| ***Tm-1 VC554 1st*** | 561 | VVKEVAVLAGVCATDPFRRMDNFLKQLESVGFCGVQNFPTVGLFDGNFRQNLEETGMGYGLEVEMIAAAH |
| ***tm-1 GCR26*** | 561 | VVKEVAVLAGVCATDPFRRMDNFLKQLESVGFCGVQNFPTVGLFDGNFRQNLEETGMGYGLEVEMIATAH |
| ***tm-1 Moneymaker*** | 561 | VVKEVAVLAGVCATDPFRRMDNFLKQLESVGFCGVQNFPTVGLFDGNFRQNLEETGMGYGLEVEMIATAH |
| ***tm-1 VC532*** | 561 | VVKEVAVLAGVCATDPFRRMDNFLKQLESVGFCGVQNFPTVGLFDGNFRQNLEETGMGYGLEVEMIATAH |
| ***Tm-1 VC554 2nd*** | 561 | VVKEVAVLAGVCATDPFRRMDNFLKQLESVGFCGVQNFPTVGLFDGNFRQNLEETGMGYGLEVEMIATAH |
| ***Tm-1 GCR237*** | 631 | RMGLLTTPYAFCPDEAVAMAEAGADIIVAHMGLTTSGSIGAKTAVSLEESVTCVQAIADATHRIYPDAIV |
| ***Tm-1 VC554 1st*** | 631 | RMGLLTTPYAFCPDEAVAMAEAGADIIVAHMGLTTSGSIGAKTAVSLEESVTCVQAIADATHRIYPDAIV |
| ***tm-1 GCR26*** | 631 | RMGLLTTPYAFCPDEAVAMAEAGADIIVAHMGLTTSGSIGAKTAVSLEESVTCVQAIADATHRINPDAIV |
| ***tm-1 Moneymaker*** | 631 | RMGLLTTPYAFCPDEAVAMAEAGADIIVAHMGLTTSGSIGAKTAVSLEESVTCVQAIADATHRINPDAIV |
| ***tm-1 VC532*** | 631 | RMGLLTTPYAFCPDEAVAMAEAGADIIVAHMGLTTSGSIGAKTAVSLEESVTCVQAIADATHRINPDAIV |
| ***Tm-1 VC554 2nd*** | 631 | RMGLLTTPYAFCPDEAVAMAEAGADIIVAHMGLTTSGSIGAKTAVSLEESVTCVQAIADATHRINPDAIV |
| ***Tm-1 GCR237*** | 701 | LCHGGPISSPEEAAYVLKRTTGVHGFYGASSMERLPVEQAITATVQQYKSISME- |
| ***Tm-1 VC554 1st*** | 701 | LCHGGPISSPEEAAYVLKRTTGVHGFYGASSMERLPVEQAITATVQQYKSISME- |
| ***tm-1 GCR26*** | 701 | LCHGGPISSPEEAAYVLKRTTGVHGFYGASSMERLPVEQAITATVQQYKSISME- |
| ***tm-1 Moneymaker*** | 701 | LCHGGPISSPEEAAYVLKRTTGVHGFYGASSMERLPVEQAITATVQQYKSISME- |
| ***tm-1 VC532*** | 701 | LCHGGPISSPEEAAYVLKRTTGVHGFYGASSMERLPVEQAITATVQQYKSISME- |
| ***Tm-1 VC554 2nd*** | 701 | LCHGGPISSPEEAAYVLKRTTGVHGFYGASSMERLPVEQAITATVQQYKSISME- |
